# Root meristem growth factor (RGF) peptide signaling as a molecular bridge between root development and non-lethal thermal stress adaptation

**DOI:** 10.1101/2025.11.27.690926

**Authors:** Yu-Chun Hsiao, Joon-Keat Lai, Shiau-Yu Shiue, Masashi Yamada

## Abstract

- Roots adapt to temperature ranges that restrict growth but are not lethal. Although lethal heat shock and moderately high temperatures have been studied in detail, the effects of non-lethal high temperatures on root development remain largely unknown. We defined 31°C as a non-lethal thermal stress in *Arabidopsis thaliana* and examined its impact on root growth using phenotypic analyses and developmental-zone-specific transcriptomics.
- Compared to growth at 22°C, at 31°C, primary root growth, meristem size, and superoxide (O₂^−^) accumulation were reduced, and the distribution of the meristem master regulator PLETHORA2 (PLT2) became restricted. Transcriptome analysis revealed a strong downregulation of *RGFs*, *RGFRs*, and *PLT2,* rather than activation of heat shock-inducible genes.
- These gene mutants were more sensitive to non-lethal thermal stress. In contrast, RGF treatment recovered heat-stress-induced defects. Beyond alleviating the stress in the primary root meristem, RGF treatments promoted lateral root elongation under prolonged non-lethal thermal stress, resulting in a more complex root system.
- These results indicate that the RGF-RGF receptor-PLT2 pathway plays a central role in root adaptation to non-lethal heat stress rather than the canonical heat shock response pathway and suggest that manipulating RGF signaling could enhance root thermotolerance and crop resilience under elevated temperatures.

## Introduction

High temperatures induce a range of developmental, physiological, and biochemical changes in plants, affecting plant growth and causing significant reductions in agricultural yields (Lobell *et al*., 2011; Lesk *et al*., 2016). Young seedlings are susceptible to high-temperature stress, which can cause significant damage to root growth and development, preventing them from accessing soil water and nutrients. Enhancing tolerance to high-temperature stress in young seedling roots improves both survival rates and yields (Lu *et al*., 2022).

In *Arabidopsis* seedlings, root developmental stages, called zonation, are well characterized. Roots exhibit dynamic growth, tightly coordinated by root zonation, which is divided into the meristematic, elongation, and differentiation zones (Petricka *et al*., 2012). Stem cell daughters undergo active cell divisions in the meristematic zone to produce numerous cells; after that, cells terminate their divisions, rapidly expand in the elongation zone, and finally differentiate to acquire distinct structures in the differentiation zone (Petricka *et al*., 2012). The size of each zone is regulated by environmental signals, phytohormones, small peptides, and endogenous cues such as reactive oxygen species (ROS) (Petricka *et al*., 2012; Hsiao & Yamada, 2020; Mase & Tsukagoshi, 2021; Hsiao & Yamada, 2024). Because the root meristem sustains continuous growth, it is highly sensitive to stress. Stress-induced reductions in meristem size compromise root function and survival (Ubogoeva *et al*., 2021), suggesting that seedlings with larger meristems may be more resilient under adverse conditions.

ROS such as superoxide (O₂^−^) and hydrogen peroxide (H₂O₂), were long regarded as harmful byproducts of metabolic stress because their excessive accumulation, particularly under stress conditions, can damage tissues and trigger cell death (Mittler, 2017). However, recent studies have shown that ROS localize to distinct root developmental zones under normal growth conditions, and their spatial distribution regulates the transition from cell division to elongation. These results indicate that ROS are key signaling molecules in root development (Dunand *et al*., 2007; Tsukagoshi *et al*., 2010; Mittler, 2017; Yamada *et al*., 2020).

As another key signaling molecule modulating root development, the ROOT MERISTEM GROWTH FACTOR 1, 2, 3 (RGF, also known as GOLVEN or CLE-like) peptides promote root meristem activity by increasing the number of meristematic cells. These three RGFs are functionally redundant, and a *rgf1/2/3* triple mutant shows a smaller root meristem phenotype. This phenotype is rescued by exogenous RGF1 treatment (Matsuzaki *et al*., 2010; Meng *et al*., 2012; Whitford *et al*., 2012). The RGF1 peptide is received by the RGF1 RECEPTORs (RGFRs)/RGF1 INSENSITIVEs (RGIs), which are specifically expressed in the meristematic zone (Ou *et al*., 2016; Shinohara *et al*., 2016; Song *et al*., 2016)

We previously showed that exogenous RGF1 elevates O₂^−^ levels in the meristematic zone while reducing H₂O₂ in the elongation and differentiation zones in the wild type, however, not in the *rgfr1/2/3* mutant (Yamada *et al*., 2020). The root meristem master regulator *PLT2* transcript is specifically expressed in the apical root meristem around quiescent center (QC) cells (Galinha *et al*., 2007). In contrast, the gradient signals of the PLT2 protein in the translational fusion line (*gPLT2-YFP*) are detected at high levels around the apical root meristem and gradually decrease until the end of the meristematic zone (Galinha *et al*., 2007). The RGF1-dependent alteration in ROS distribution post-translationally stabilizes the PLT2 protein and enlarges the meristem size by broader PLT2 gradients (Yamada *et al*., 2020; Hsiao *et al*., 2025). These findings demonstrate that RGF-RGFR-PLT2 signaling regulates meristem size via ROS (Yamada *et al*., 2020; Hsiao *et al*., 2025).

In addition to RGF1-3, other RGF members, such as RGF5 and RGF8, are involved in lateral root development (Fernandez *et al*., 2013). The *rgf5/8* double mutant increases total lateral root primordium density as compared with the single mutants and the wild type (Fernandez *et al*., 2020). Overexpression of *RGF5* and *RGF8* and treatment with high concentrations of exogenous RGF5 and RGF8 peptides inhibit the formation of the lateral root primordium (Fernandez *et al*., 2015; Fernandez *et al*., 2020). The lateral root development in the triple mutant background of the *RGFR1/RGI1*, *RGI4*, and *RGI5* receptor genes, which is not inhibited by *RGF5* overexpression, suggests *RGF5*/*RGF8*-*RGI* receptor pathways modulate lateral root development (Fernandez *et al*., 2020).

As an environmental signal, high temperatures elicit distinct root responses depending on intensity. Acute heat shock (40-42°C) is lethal to *Arabidopsis* seedlings, but prior exposure to sublethal levels (37°C) induces acclimation through the accumulation of heat shock proteins and ROS-scavenging enzymes (Yeh *et al*., 2012; Ohama *et al*., 2017). By contrast, moderately high temperatures (26-29°C) promote root growth via hormone-mediated thermomorphogenesis (Quint *et al*., 2016; Vu *et al*., 2019). We can expect that non-lethal thermal stress above 30°C strongly suppresses root growth, limits water and nutrient availability, and ultimately affects whole-plant development. It is essential to understand the mechanisms underlying acclimation to non-lethal thermal stress and the benefits of this process for improving thermal resilience. However, it remains unclear.

In this study, we define 31°C as a non-lethal yet growth-restrictive temperature that limits root meristem development. Transcriptome profiling reveals strong repression of *RGF2*, *RGFR2*, and *PLT2* instead of induction of the canonical stress response genes, and mutants lacking these genes are particularly sensitive in root meristem and lateral root development to non-lethal thermal stress. Exogenous RGF application counteracts these inhibitory effects of root meristem development and lateral root growth. Together, these findings demonstrate that RGF peptide signaling plays a crucial role in root adaptation to non-lethal thermal stress by modulating established root developmental programs rather than stimulating the canonical heat-shock responses at elevated temperatures. They also highlight the potential of RGF peptide application as a strategy to counteract heat-induced root growth inhibition and to strengthen the resilience of root architecture in warming environments.

## Materials and Methods

### Plant material and growth conditions

All *Arabidopsis* mutants and transgenic lines used in this study (SALK_130119.20.25 (*plt2*), *rgf123*, *rgfr123, gPLT2-YFP*) are in the Columbia-0 (Col-0) genetic background and have been previously characterized (Galinha *et al*., 2007; Matsuzaki *et al*., 2010; Shinohara *et al*., 2016; Yamada *et al*., 2020; Hsiao *et al*., 2025). Seeds were surface-sterilized with 50% bleach supplemented with 0.1% Tween 20 (Sigma) for 10 minutes, followed by five rinses with sterile distilled water. The sterilized seeds were then stratified at 4°C in the dark for 2 days. Subsequently, seeds were sown on half-strength Murashige and Skoog (½ MS) medium (Caisson Laboratories) supplemented with 0.05% MES, 1% sucrose, and 1% agar (Sigma) and pH adjusted to 5.7 with KOH. Seedlings were grown vertically in a growth chamber (F-600 HiPoint, Taiwan) at 22°C under long-day conditions (16 h light/8 h dark) for 7 days. After this period, seedlings were transferred to ½ MS agar plates containing either sterile water (mock treatment), synthetic sulfated RGF1 peptide (Mission Biotech, Taiwan), or subjected to the indicated temperature treatments for 1 to 3 days for further experiments.

### Root growth assay

Seven-day-old *Arabidopsis* seedlings were transferred to growth chambers set at different temperatures (27-32°C) for 3 days. Primary root growth was observed and recorded daily at a consistent time, with a mark applied to the root tip each day. After the final observation on day three, images of the plates were scanned using an Epson Perfection V800, and root lengths were measured using Fiji ImageJ (Schindelin *et al*., 2012) and normalized relative to those of seedlings grown at 22°C.

For lateral root observation, seven-day-old *Arabidopsis* seedlings were transferred to the indicated media and grown at 22°C or 31°C for six days. Root images were captured using the same scanner, and lateral root lengths were measured with Fiji ImageJ (Schindelin *et al*., 2012). The average lateral root length per seedling was calculated and used for statistical analysis.

### Light and confocal microscopy analysis

For superoxide (O_₂_^−^) detection, seedlings were stained with 400 *μ*M nitroblue tetrazolium (NBT) dissolved in 20 mM phosphate buffer (pH 6.1) in the dark for 2 minutes, followed by two rinses with distilled water, as previously described (Hsiao *et al*., 2025). Root images were captured using a 10× objective lens on a Zeiss AXIO Scope A1 microscope.

Root tip structures and fluorescent signals were observed using a 20× objective on a Zeiss LSM 980 laser scanning confocal microscope. Excitation and emission settings were as follows: YFP, excitation at 514 nm and emission at 517-544 nm; propidium iodide staining, excitation at 561 nm and emission at 565-747 nm. Confocal images were stitched and analyzed using Fiji ImageJ (Schindelin *et al*., 2012), and stitched mosaics were assembled (Thevenaz & Unser, 2007). Each confocal image was stitched and overlaid on a black background to enhance visual consistency and improve readability. The meristematic zone was defined as the region extending from the apical root meristem up to the cortex cell whose longitudinal (y-axis) length was more than twice that of the preceding cell, and the number of cortex cells within this region was counted to estimate meristem size.

### RNA-seq library preparation and data processing

Similarly sized root tip tissues, including the three developmental zones, were precisely cut on a 2% agar plate containing ½ MS using an ophthalmic scalpel (Feather) under a dissection microscope (Leica M205 FCA), then homogenized in RNA extraction buffer, and processed according to the manufacturer’s recommended protocol. RNA extracted from 50 root tissues was pooled for each sample. Total RNA was extracted using the RNeasy Micro Kit (Qiagen) according to the manufacturer’s instructions. RNA concentration was quantified using a Qubit fluorometer (Invitrogen). RNA quality was assessed using a 2100 Bioanalyzer (Agilent), and all samples exhibited RNA integrity numbers above 8.0. RNA-seq libraries were prepared from 300 ng of total RNA using the Universal Plus Total RNA-Seq with NuQuant Library Preparation Kit (Tecan Trading AG), following the manufacturer’s instructions. Libraries from 5 biological replicates of root samples under the control temperature (22°C) and non-lethal high temperature (31°C) at 1 and 2 days after transfer (DAT) were sequenced on an Illumina NextSeq2000 platform, generating 150-base paired-end reads. The RNA-sequencing reads were aligned to the *Arabidopsis* reference genome TAIR10 using subread (Liao *et al*., 2013). The mRNA read counts of genes were computed using featureCounts (Liao *et al*., 2014). Genes expressed (read counts > 0) are included for analysis. Normalization for sequencing depth and differential gene analysis was performed using DESeq2 (Love *et al*., 2014). Genes with FDR < 5% were considered differentially expressed. GO analysis was performed using the Bioconductor package clusterProfiler (Xu *et al*., 2024). All DEGs can be found in Supporting Information Dataset S1.

### Statistical analysis and reproducibility

Experiments were independently repeated 3 times with similar results. No power analysis was done to estimate the sample size. All statistical analyses were performed using R version 4.5.0 (http://www.r-project.org/). For 2-sample comparison, data were analyzed with a 2-tailed t-test or Wilcoxon test depending on the data normality. As an omnibus test, an analysis of variance (ANOVA) or an aligned rank transform (ART)-ANOVA was performed, depending on the data normality. As a post hoc test, ART-Contrast test was applied to non-normal data, and the Games–Howell test, Tukey’s tests, or REGWQ test was performed on normal data, depending on the data homoscedasticity and sample sizes. In the boxplots, the circles represent each individual data point; the horizontal line within boxes represents the median; the upper and lower hinges, respectively, indicate the 75^th^ and 25^th^ percentiles (interquartile range [IQR]); and the whiskers show the 1.5 extension of the IQR.

### Data availability

The RNA-seq data generated in this study will be deposited in a public repository (e.g., NCBI GEO or an equivalent platform) once access to deposition services is restored. All other data supporting the findings of this study are included in the article and its supplementary materials, and additional information is available from the corresponding author upon reasonable request.

## Results

### Temperatures between 30-32°C inhibit root growth, reduce meristem size, and suppress O₂^−^ accumulation

Sublethal but elevated temperatures inhibit root growth; however, this effect remains uncharacterized. We first wished to identify the threshold temperature at which root growth is impaired. To do so, *Arabidopsis* seedlings were grown at 22°C for 7 days and then transferred to either the control condition (22°C) or elevated temperatures (27, 28, 29, 30, 31, or 32°C). Root length was measured every 24 h for 3 days, and growth ratios were calculated relative to 22°C. At 1 DAT (1 day after transfer), seedlings transferred to 32°C showed reduced root growth, while the growth ratios of seedlings transferred to other temperatures were not statistically different from those maintained at 22°C (Fig. 1a, b). By 2 DAT, however, growth ratios of plants transferred to 30°C, 31°C, and 32°C declined. By 3 DAT root growth was modestly reduced in seedlings transferred to 29°C, strongly suppressed at 30°C and 31°C, and nearly abolished at 32°C (Fig. 1a, b). These results demonstrate that temperatures of 30-32°C adversely affect root growth.

**Fig. 1.**
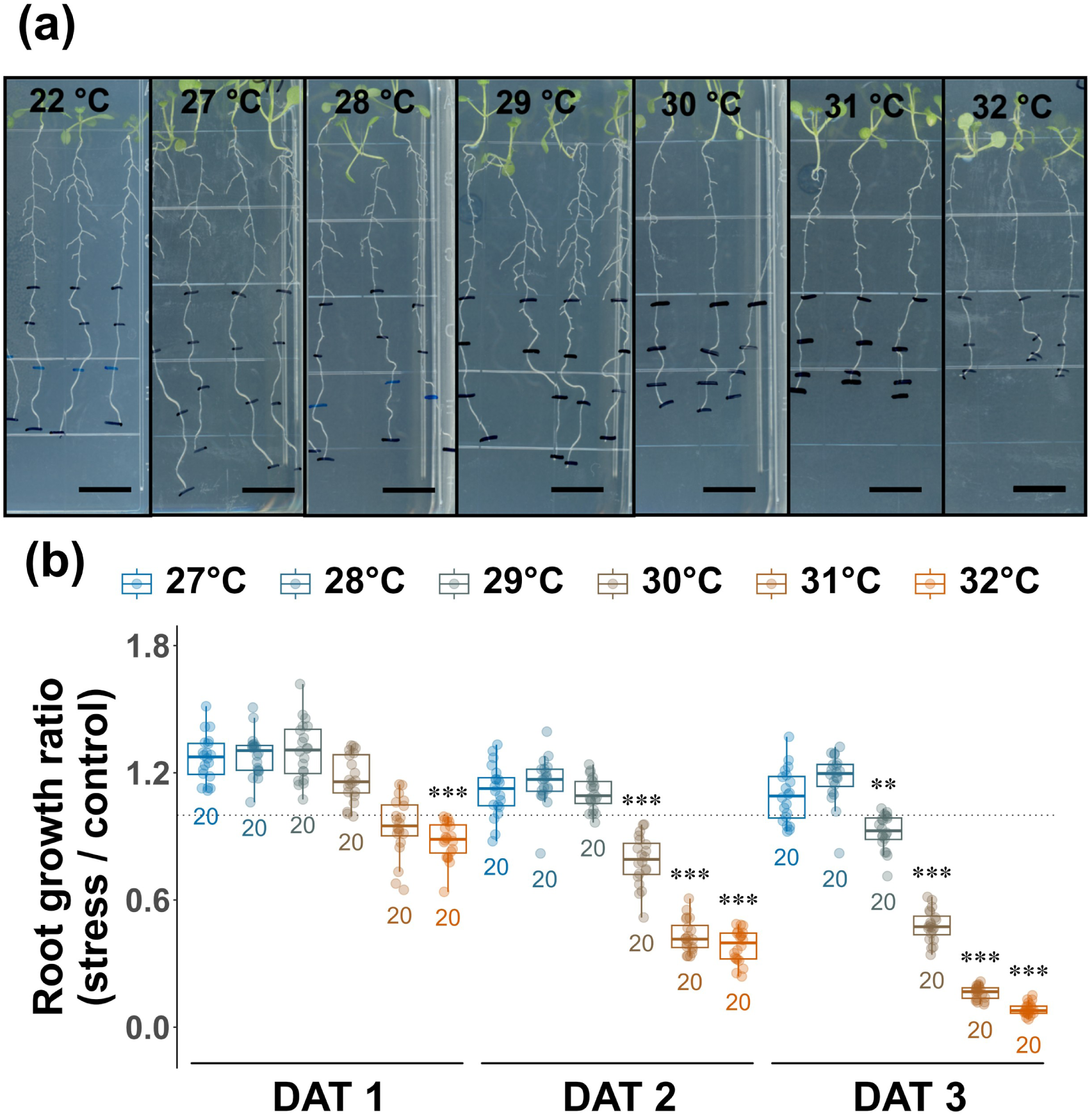
High temperature inhibits root growth. (a) Primary root lengths were recorded daily for 3 days after transfer (DAT) to control (22°C) or to elevated temperatures (27-32°C). The topmost black line in each image indicates the root tip position at 0 DAT marked on the petri dish lid. The locations of root tips were subsequently marked every 24 h. Scale bar = 10 mm. (b) Root growth ratios were calculated as the relative root length under thermal stress (27-32°C) compared with the control condition (stress/control) at each of the 3 DAT. The average root length measured at 22°C was defined as 1. Asterisks indicate where the root growth ratio of the heat treatment group is significantly less than the control (t-test or Wilcoxon rank sum test, depending on the data normality; ***, p < 0.001; **, p < 0.01; *, p < 0.05). Integer values below the boxplots represent the sample size of each group. Grey-dotted line marks the ratio value of 1.

Spatial distributions of ROS in the root developmental zones modulate root growth (Dunand *et al*., 2007; Tsukagoshi *et al*., 2010). O₂^−^ specifically accumulates in the meristematic zone, in correlation with root length and meristematic zone size (Tsukagoshi *et al*., 2010; Yamada *et al*., 2020). We therefore hypothesized that O₂^−^decreases concomitantly with root growth inhibition under elevated temperatures. To test this, O₂^−^ levels were assessed by nitro blue tetrazolium (NBT) staining. Staining intensity in the meristematic zone decreased in seedlings transferred to warmer temperatures at 2 DAT and was markedly reduced by 3 DAT at 31-32°C compared with 22°C (Fig. 2a, b). Thus, root growth inhibition (Fig. 1) and reduced O₂^−^accumulation (Fig. 2) occurred in parallel at 31-32°C. We noted that, while root growth was almost completely arrested at 32°C, roots at 31°C continued to elongate slowly (Fig. 1a), indicating that 31°C provides a non-lethal condition suitable for studying root adaptation. Concurrent reductions of root growth and O₂^−^ accumulation suggest downsizing of the meristematic zone under non-lethal thermal stress (31°C). To test this, we measured cell number in the meristematic zone as tracked meristem zone size using confocal imaging under non-lethal thermal stress at the same time points. We found that at 1 DAT, meristematic zone size was similar in plants that had been transferred to 31°C and 22°C (Fig. 3a, b). By 2-3 DAT, however, the cell number in the meristematic zone was significantly reduced at 31°C (Fig. 3a, b), consistent with the reduced root growth and O₂^−^ accumulation we observed in Figs 1-2. Despite this reduction of meristematic zone size, the apical root meristem retained its structural integrity without any typical damage to the apical root meristem, such as columella and QC cell disruptions (Fig. 3a). Instead of these damages, premature cell elongation and differentiation were observed at 31°C compared with 22°C (Fig. 3a, arrowheads show the junction between the meristematic zone and elongation zone). Together, these findings show that non-lethal thermal stress of 31°C restricts the meristematic zone and accelerates cell differentiation through a coordinated mechanism. Having established 31°C as a non-lethal thermal stress condition with key characteristics of meristem growth restriction, we next examined the molecular basis of root adaptation under non-lethal thermal stress condition (31°C).

**Fig. 2.**
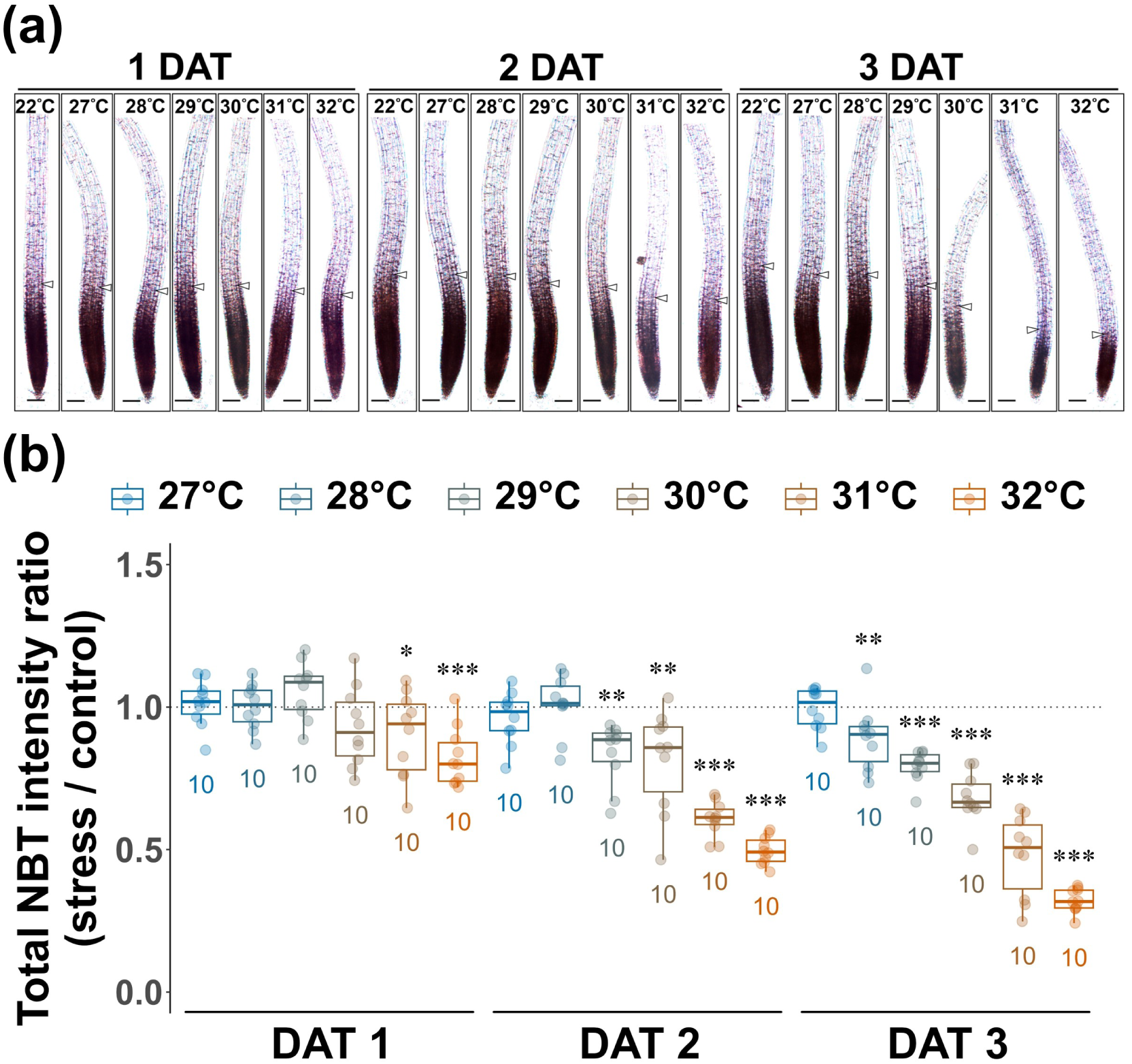
High temperature stress decreases O_2_^−^ levels in the root meristematic zone. (a) Light microscopy images of Col-0 root tips stained with nitro-blue tetrazolium (NBT) at 1, 2, or 3 days after transfer (DAT) to either control conditions (22°C) or elevated temperatures (27-32°C). Images were captured under bright-field illumination at 10x magnification. Representative pictures shown; n = 3 independent experiments were performed. Each experiment included more than 15 roots. Arrowheads indicate the junction between the meristematic and elongation zones (see methods). Scale bar = 100 µm. (b) Quantitative analysis of NBT staining in Col-0 roots under the indicated conditions. The intensity of the control group (22°C) was set to 1. Asterisks indicate that the NBT total intensity ratio of the heat treatment group is significantly less than the control (t-test or Wilcoxon rank sum test, depending on the data normality; ***, p < 0.001; **, p < 0.01; *, p < 0.05). The integer values below the boxplots represent the sample sizes of each group. The grey-dotted line marks the ratio value of 1.

**Fig. 3.**
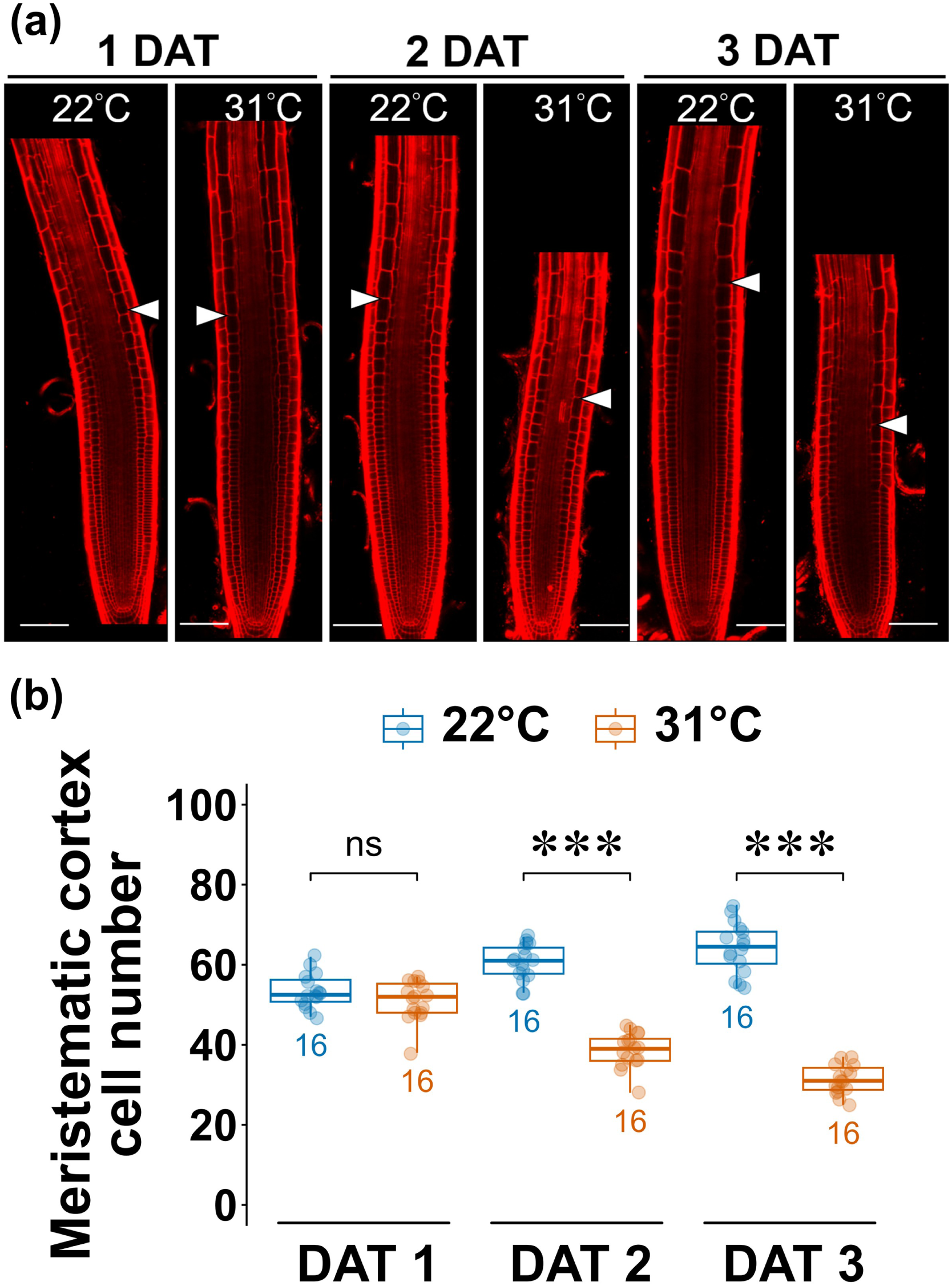
The size of the meristematic zone decreases during adaptation to high heat stress at 31°C. (a) Confocal microscopy was used to capture daily images of the three developmental zones for 3 days after transfer (DAT) to control (22°C) and heat-stress temperature (31°C) conditions. Arrowheads indicate the boundary between the meristematic zone and the elongation zone, defined as the region between the quiescent centre and the last cortical cell that had not doubled in length relative to its predecessor. Representative images are shown; n = 3 independent experiments. Each experiment included more than 15 roots. Scale bar = 100 µm. (b) Quantitative analysis of meristematic cortex cell number in roots grown under the indicated conditions. Asterisks indicate that the meristematic cortex cell of the heat treatment group is significantly different from the control (t-test; ***, p < 0.001; **, p < 0.01; *, p < 0.05), while ns indicates non-significant differences (p > 0.05). In the boxplots, the circles represent each individual data point, and sample sizes are shown below the corresponding boxes. The horizontal line within the boxes represents the median, and the upper and lower hinges, respectively, indicate the 75th and 25th percentiles (interquartile range, IQR). The whiskers show the 1.5 extension of the IQR.

### Non-lethal thermal stress modulates the RGF peptide-RGF receptor-PLT2 signaling pathway

Most transcriptome studies in *Arabidopsis* have focused on whole seedlings or entire roots subjected to either lethal or moderately high temperatures (Larkindale & Vierling, 2008; Quint *et al*., 2016; Vu *et al*., 2019; Guo *et al*., 2022). While such studies have provided important global insights, they lack spatial resolution and do not address non-lethal thermal stress. More recently, single-cell RNA-Seq under lethal heat stress (Jean-Baptiste *et al*., 2019) has enabled cell-type-specific profiling, but zone-specific responses to non-lethal temperatures remain poorly understood.

To address this gap, we performed transcriptome analysis of three root developmental zones under non-lethal thermal stress (31°C). Total RNA was collected from seedlings 1 and 2 DAT to 22°C or 31°C, and comparative analyses were conducted. We identified more than 1,200 differentially expressed genes (DEGs) at 1 DAT, over 2,600 at 2 DAT, and approximately 4,200 across both time points (Fig. 4a). In agreement with the phenotypic observation that meristematic zone size was reduced only at 2 DAT (Fig. 3), gene ontology (GO) enrichment of down-regulated genes at 2 DAT revealed significant associations with meristem development, the mitotic cell cycle, and DNA replication (Fig. 4b).

**Fig. 4.**
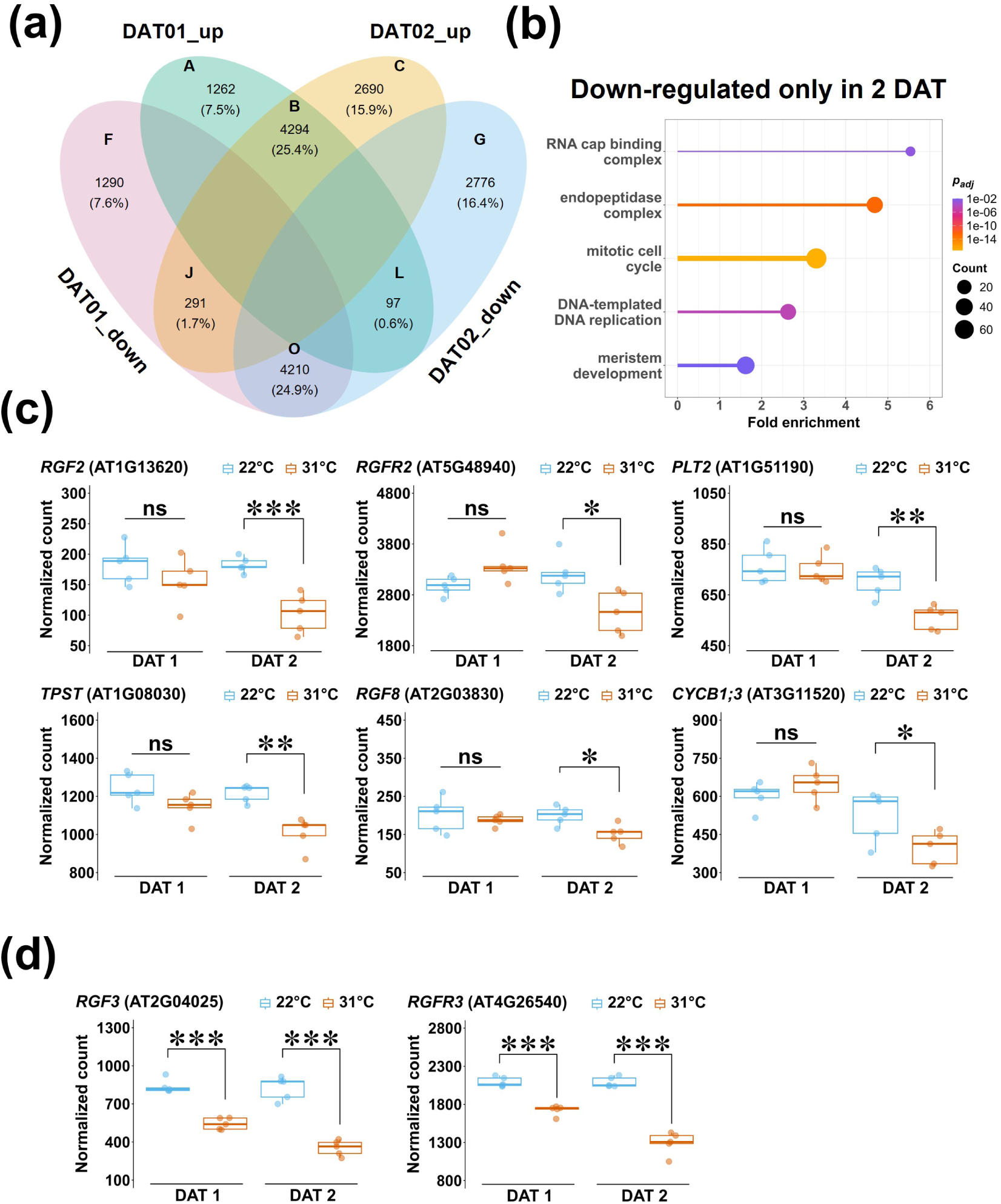
High thermal stress diminishes the expression of genes related to the RGF1-receptor-PLT2 pathway and stimulates ROS metabolic processes and ROS response-relevant genes. (a) Four-set Venn diagram showing the number of differentially expressing genes (DEGs) under heat-stress temperature (31°C) compared to the control (22°C) condition. Up-regulated genes in 1 day after transfer (DAT) and/or 2 DAT were respectively encompassed in the green and orange ellipses. Down-regulated genes in 1 DAT and/or 2 DAT were respectively encompassed in the red and blue ellipses. Each subset of DEGs were named with a capital letter. The integer value represents the number of DEGs in each subset. The percentage represents the proportion of the DEGs number in each set from the total DEGs. (b) Gene ontology (GO) analysis of down-regulated genes after 2 days of non-lethal heat treatment (subset G). (c) RGF1-receptor-PLT2 pathway related genes from subset G (down-regulated only in 1 DAT). (d) RGF1-receptor-PLT2 pathway related genes from subset O (down-regulated in 1 and 2 DAT). Asterisks indicate the gene expression of the heat treatment group is significantly different from the control (t-test; ***, p < 0.001; **, p < 0.01; *, p < 0.05), while *ns* indicates non-significant differences (p > 0.05). In the boxplots, the circles represent each individual data point, and sample sizes are shown below the corresponding boxes. The horizontal line within the boxes represents the median, and the upper and lower hinges, respectively, indicate the 75^th^ and 25^th^ percentiles (interquartile range, IQR). The whiskers show the 1.5 extension of the IQR.

Notably, in these GOs, genes central to the RGF peptide signaling cascade, including *TYROSYLPROTEIN SULFOTRANSFERASE* (*TPST*), *RGF2*, *RGF1 RECEPTOR2* (*RGFR2*), and *PLT2*, were significantly downregulated at 2 DAT at 31°C, but not at 1 DAT (Fig. 4c). The *RGF1/2/3* and *RGFR1/2/3* genes function redundantly in root meristem development (Matsuzaki *et al*., 2010; Shinohara *et al*., 2016). Additionally, we examined expression of these genes and found that *RGF3* and *RGFR3* began to decrease at 1 DAT and further decreased by 2 DAT (Fig. 4d). These results indicate that non-lethal thermal stress suppressed the expression of key components of the *RGF*-*RGFR*-*PLT2* pathway at the time when meristem reduction became apparent.

These results suggested that roots adapt to non-lethal thermal stress by modulating genes related to root development rather than canonical stress response genes. To directly test this idea, we used *HSFA2*, *HSFA7a*, *HSP70-1*, and *HSP101* as representative molecular markers of the heat-shock response (Schramm *et al*., 2006; Charng *et al*., 2007; Liu *et al*., 2011; Yoshida *et al*., 2011). These genes are known downstream targets of *HSFA1*s, the master regulators of acquired thermotolerance, and are strongly induced during acute heat shock. Consistent with this, when we compared the root transcriptome (22 °C and 31 °C for 1 and 2 DAT across three developmental zones) with a published heat-shock dataset (22°C and 44°C for 3 h in 7-day-old seedlings) (Yang *et al*., 2023), the expression levels of *HSFA2*, *HSFA7a*, *HSP70-1*, and *HSP101* were highly upregulated under heat shock but showed only modest or negligible induction under non-lethal thermal stress (Fig. S1).

Together, these results indicate that non-lethal heat stress triggers a regulatory program distinct from acute heat shock, promoting developmental adjustment rather than activating the canonical *HSFA1s*-*HSP* signaling cascade. Roots, therefore, appear to adapt to non-lethal thermal stress mainly by modulating developmental gene networks rather than by strongly inducing classical stress-responsive genes.

### The *rgf1/2/3*, *rgfr1/2/3*, and *plt2* mutants are hypersensitive to non-lethal thermal stress

Together with the phenotypic data shown in Figs 1–3, our transcriptomic findings suggested that the RGF peptide-RGF receptor-PLT2 signaling pathway controls root meristem size during adaptation to non-lethal thermal stress. To test this idea, we examined meristem size and O₂^−^ accumulation in *rgf1/2/3*, *rgfr1/2/3*, and *plt2* mutants exposed to non-lethal thermal stress.

In wild-type seedlings, the meristematic zone size at 31°C was comparable to that at 22°C at 1 DAT, but significantly reduced by 2 DAT (Fig. 5a, c). In contrast, in both *rgf1/2/3* and *rgfr1/2/3* mutants, the relative meristematic zone size at 31°C compared with that at 22°C was smaller already at 1 DAT, with further reductions by 2 DAT (Fig. 5a, c). A similar trend was observed for O₂^−^ levels: in the wild type, NBT staining at 31°C was indistinguishable from that at 22°C at 1 DAT, but decreased by 2 DAT (Fig. 5b, d). However, in the *rgf1/2/3* and *rgfr1/2/3* mutants, O₂^−^ signals under non-lethal thermal stress compared with the control temperature showed weaker O₂^−^ signals as early as 1 DAT, which became strongly diminished by 2 DAT (Fig. 5b, d). Similarly, in the *plt2* mutant, the relative meristematic zone size and O₂^−^ accumulation under non-lethal thermal stress compared with the control temperature were smaller at both 1 and 2 DAT compared with the wild type (Fig. 5a–d). Together, these findings demonstrated that the *rgf1/2/3*, *rgfr1/2/3,* and *plt2* mutants showed hypersensitive phenotypes in meristematic zone size and O₂^−^ levels under non-lethal thermal stress compared with the control temperature, indicating that thermal stress reduces meristem size and O₂^−^ levels via the *RGF*-*RGFR*-*PLT2* signaling pathway.

**Fig. 5.**
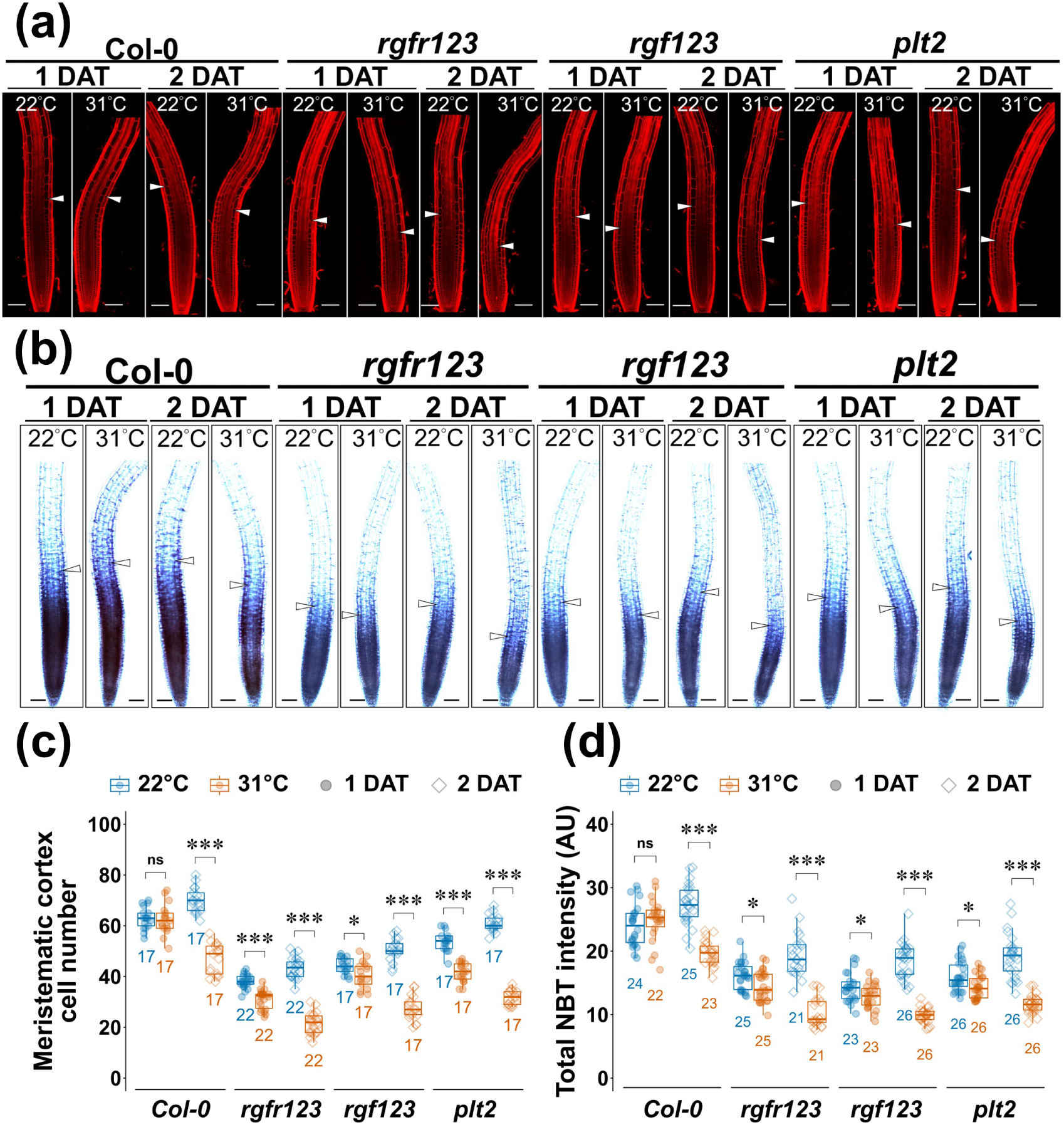
The *rgf1/2/3*, *rgfr1/2/3*, and *plt2* mutants exhibit a phenotype that is sensitive to thermal stress. (a) Confocal images of roots from wild-type (Col-0) and mutant lines at 1 or 2 days DAT to control (22°C) or heat-stress (31°C) conditions. Roots were stained with propidium iodide before imaging. (b) Light microscope images of roots from designated genotypes at 1 or 2 DAT to control (22°C) or heat-stress (31°C) conditions, followed by nitro-blue tetrazolium staining. The arrowheads in (a) and (b) indicate the junction between the meristematic and elongation zones. All scale bar = 100 µm. (c–d) Quantitative analysis of (a) and (b). Asterisks indicate the heat treatment group is significantly different from the control (t-test or Wilcoxon rank sum test, depending on the data normality; ***, p < 0.001; **, p < 0.01; *, p < 0.05), while ns indicates non-significant differences (p > 0.05). In the boxplots, the circles represent each individual data point, and sample sizes are shown below the corresponding boxes. The horizontal line within the boxes represents the median, and the upper and lower hinges, respectively, indicate the 75^th^ and 25^th^ percentiles (interquartile range, IQR). The whiskers show the 1.5 extension of the IQR.

### RGF1 peptide treatment mitigates non-lethal thermal stress and restores root meristem size

The down-regulation of the *RGF*-*RGFR*-*PLT2* pathway genes (Fig. 4c, d) coincided with reduced meristem size after 2 DAT to non-lethal thermal stress (Fig. 3), suggesting that thermal stress impairs root meristem development by inhibiting this pathway. We therefore hypothesized that exogenous RGF1 application could counteract these defects by activating the RGF-RGFR-PLT2 pathway at 2 DAT, when thermal stress defects are initiated.

To test this hypothesis, we grew the seedlings of the *gPLT2-YFP* reporter line (Galinha *et al*., 2007) for 7 days at 22°C, transferred them to 22°C or 31°C, and applied mock or 5 nM RGF1 peptide treatment at 2 DAT at 31°C. At 3 DAT, we assessed meristem size and O₂ accumulation. To monitor whether RGF1 activates the RGF-RGFR-PLT2 pathway, we also assessed PLT2-YFP protein localization. In mock-treated seedlings at 31°C, meristem size was reduced compared with controls at 22°C (Fig. 6a, d). However, RGF1-treated seedlings showed a significantly larger meristem, even under the heat stress (Fig. 6a, d). Consistently, O₂^−^ accumulation was higher with RGF1 treatment than with mock treatment at 31°C (Fig. 6b, e). Non-lethal thermal stress also restricted the PLT2 protein localization domain compared with control conditions (Fig. 6c, f). Strikingly, RGF1 application expanded PLT2 distribution despite the stress (Fig. 6c, f). Together, these results demonstrate that RGF1 treatment mitigates the effects of non-lethal heat stress on O₂^−^ production, meristem size, and PLT2 protein expression.

**Fig. 6.**
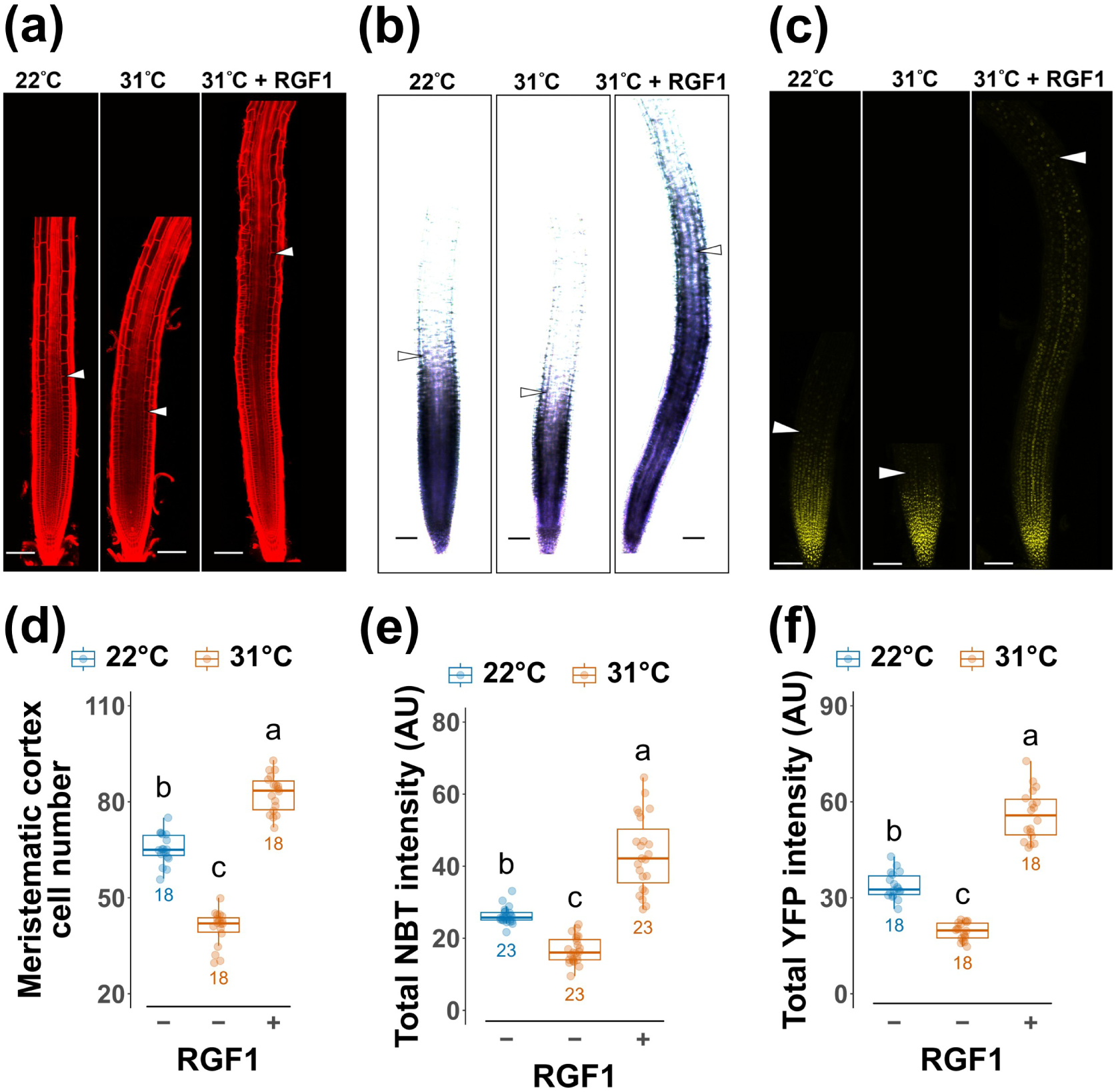
RGF1 treatment after 1 DAT to 31°C mitigates thermal stress. (a) Confocal images of gPLT2-YFP roots 2 DAT to control (22°C) or heat-stress (31°C) conditions with or without 5 nM RGF1 treatment; roots were stained with propidium iodide. (b) Light microscope images of gPLT2-YFP roots 2 DAT to 22°C or 31°C conditions with or without 5 nM RGF1 treatment; stained with nitro-blue tetrazolium. Arrowheads in (a) and (b) mark the junction between the meristematic and elongation zones. (c) Confocal images of gPLT2-YFP roots 2 DAT to the indicated conditions. The confocal images in (a) and (c) are composed of multiple stitched tiles, set against a black background. All scale bars = 100 µm. (d-f) Quantitative analysis of images in (a‒c). Different letters represent significant differences using Fisher’s ANOVA + REGWQ test (d), ART-ANOVA + ART-Contrast test (e), or Welch’s ANOVA + Games-Howell test (f), p < 0.05. In the boxplots, the circles represent each individual data point, and sample sizes are shown below the corresponding boxes. The horizontal line within boxes represents the median, and the upper and lower hinges, respectively, indicate the 75^th^ and 25^th^ percentiles (interquartile range, IQR). The whiskers show the 1.5 extension of the IQR.

### The RGF2 and RGF8 peptide treatments stimulate lateral root growth under extended heat stress

We investigated non-lethal thermal stress adaptation in the primary root meristem at the early seedling stage (7-9-day-old). Importantly, the impairments in meristem function observed under non-lethal stress were not transient. They persisted into later developmental stages (13-day-old), reducing both primary root elongation and lateral root development. We explored thermal-stress adaptation in lateral roots. We hypothesized that RGF2 and RGF8 mitigate lateral root growth inhibition during extended non-lethal thermal stress, because these two peptide genes were notably downregulated under non-lethal thermal stress in our transcriptome analysis (Fig. 4c).

To test this hypothesis, we examined whether the RGF2 and RGF8 peptides can also alleviate lateral root development inhibition during extended non-lethal thermal stress, beyond alleviating the root meristem inhibition in the primary root.

Seedlings were grown for seven days at 22°C and then transferred to either the control temperature (22°C) or the non-lethal thermal stress condition (31°C). Plants were subsequently mock-treated, or exposed to RGF2 or RGF8 for 6 days (6 DAT; 13-day-old seedlings) (Fig. 7). Roots produced numerous and elongated lateral roots under the control condition (22°C), and treatment with low concentrations (10, 50, and 100 pM) of the RGF2 or RGF8 peptides had little effect on lateral root growth (Fig. 7a, c). Exposure to 31°C with mock treatment markedly suppressed this lateral root growth (Fig. 7a, b). However, RGF2 and 8 treatments at 50 pM substantially restored lateral root development under the heat stress, alleviating the inhibition observed at 31°C (Fig. 7b, d). Interestingly, lower concentrations of peptides did not have any effect under the control condition (Fig. 7a, c) but did strongly stimulate lateral root elongation under non-lethal thermal stress (Fig. 7b, d).

**Fig. 7.**
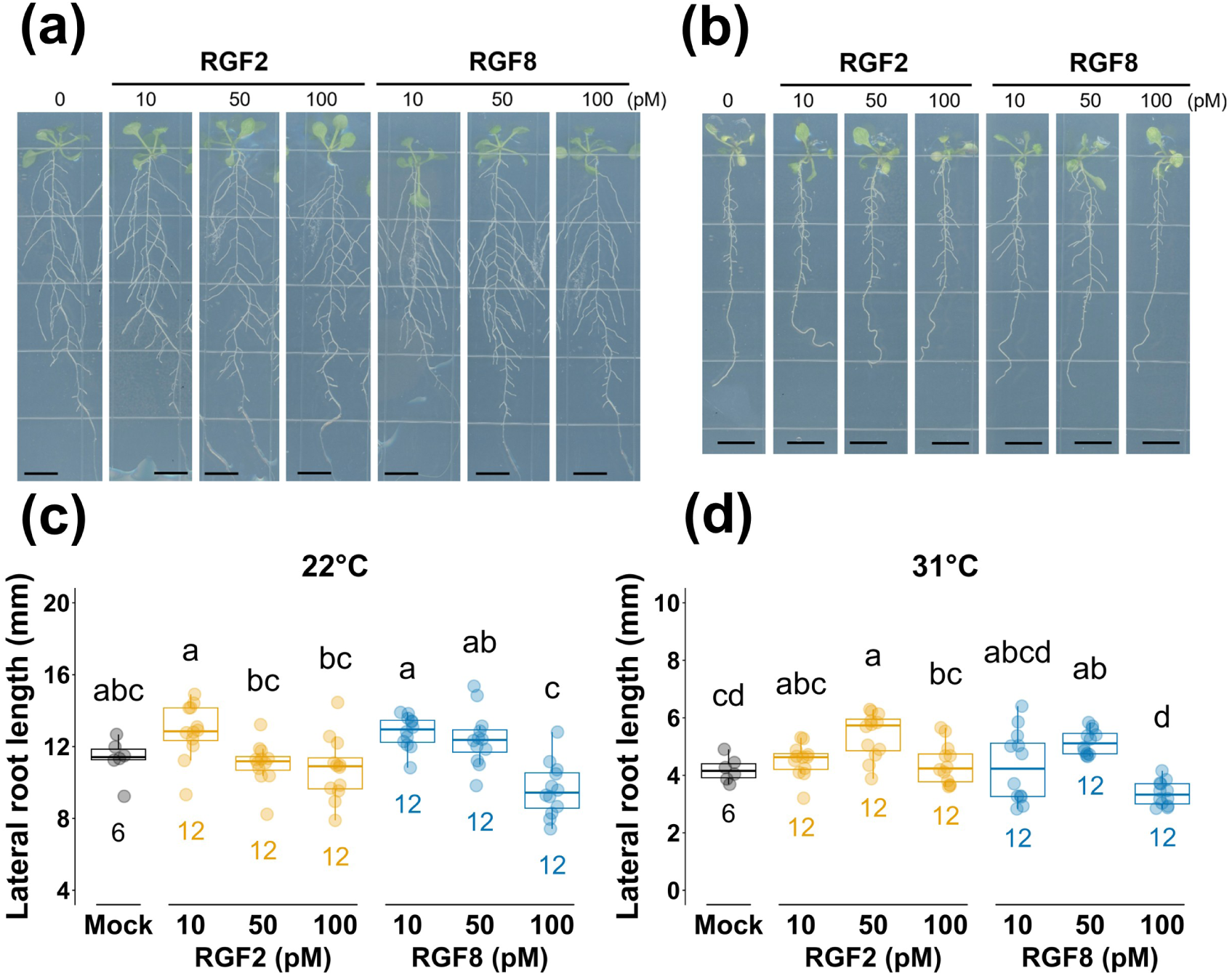
RGF2 and RGF8 peptide treatments stimulate lateral root growth under non-lethal heat stress. (a‒b) Light microscope images taken after wild-type seedlings were transferred to the indicated concentrations of RGF2 or RGF8 peptides for six days under (a) 22°C or (b) 31°C conditions. All scale bars = 10 mm. (c, d) Quantitative analysis of lateral root length in seedlings treated as shown in (a) and (b), respectively. Different letters represent significant differences using Fisher’s ANOVA + Tukey-Kramer test (c) and Welch’s ANOVA + Games-Howell test (d), p < 0.05. In the boxplots, the circles represent each individual data point, and sample sizes are shown below the corresponding boxes. The horizontal line within boxes represents the median, and the upper and lower hinges, respectively, indicate the 75^th^ and 25^th^ percentiles (interquartile range, IQR). The whiskers show the 1.5 extension of the IQR.

Taken together, these data suggest that the defects in lateral root development under non-lethal thermal stress are primarily caused by the observed downregulation of *RGF2* and *RGF8*, and that RGF2 and RGF8 treatments enhance lateral root growth under non-lethal thermal stress. Thus, RGF peptides not only sustain primary root meristem activity but also promote lateral root development, resulting in a more elaborate and potentially more resilient root system under non-lethal thermal stress.

### The *RGF*-*RGFR*-*PLT2* pathway is essential for lateral root growth

Because the *rgf1/2/3*, *rgfr1/2/3*, and *plt2* mutants exhibited hypersensitive responses to non-lethal thermal stress, showing smaller root meristems and reduced superoxide accumulation (Fig. 5), we wondered if they would show a similar hypersensitive lateral root response. To ask this question, we examined their lateral root phenotypes at 6 DAT (13-day-old seedlings) under both control (22°C) and non-lethal thermal stress (31°C) conditions (Fig. 8).

**Fig. 8.**
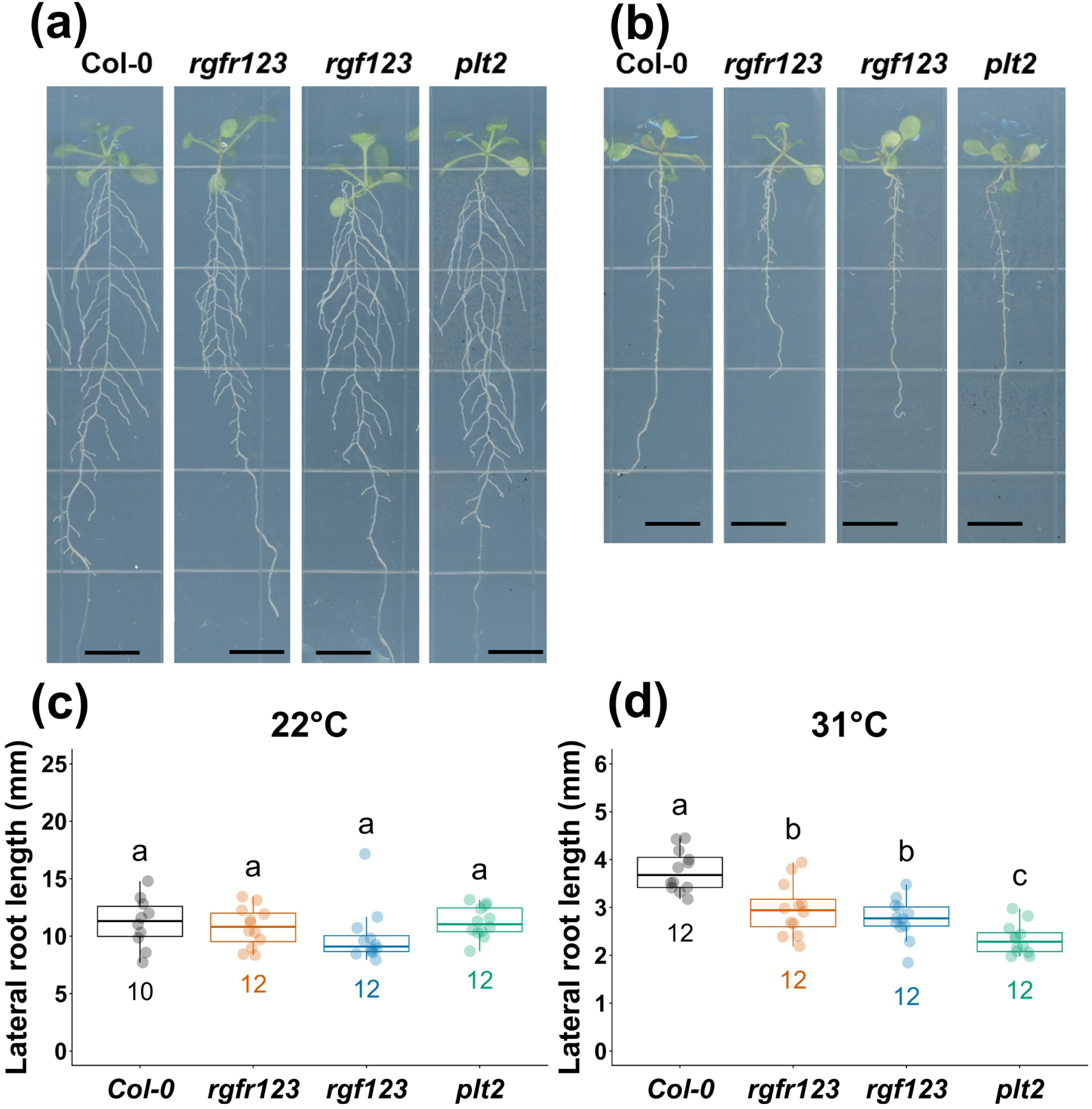
The lateral roots and leaves of the *rgf1/2/3*, *rgfr1/2/3*, and *plt2* mutants are sensitive to non-lethal heat stress. (a‒b) Images of wild-type (Col-0) and mutant lines were taken after transfer to (a) 22°C or (b) 31°C for six days. All scale bars = 10 mm. (c, d) Quantitative analysis of lateral root length in seedlings treated as in (a) and (b), respectively. Different letters represent significant differences using ART-ANOVA + Tukey-Kramer test (c) and Fisher’s ANOVA + REGWQ test (d), p < 0.05. In the boxplots, the circles represent each individual data point, and sample sizes are shown below the corresponding boxes. The horizontal line within boxes represents the median, and the upper and lower hinges, respectively, indicate the 75th and 25th percentiles (interquartile range, IQR). The whiskers show the 1.5 extension of the IQR.

At 22°C, 13-day-old *rgf1/2/3*, *rgfr1/2/3*, and *plt2* mutant seedlings developed lateral roots comparable to those of the wild type (Fig. 8a, c). However, exposure to 31°C markedly restricted lateral root elongation in the mutants, resulting in simpler root systems with shorter lateral roots by 6 DAT compared with plants grown at 22°C (Fig. 8b, d). Thus, the *rgf1/2/3*, *rgfr1/2/3*, and *plt2* mutants exhibited more pronounced defects than the wild type in both primary root and lateral roots under non-lethal thermal stress.

## Discussion

A deeper understanding of how non-lethal thermal stress reshapes root developmental programs will be essential for developing strategies to enhance crop resilience and maintain productivity under future climate scenarios.

### Defining 31°C as a non-lethal thermal stress condition

Defining non-lethal thermal stress conditions is important step in better understanding how plants can be expected to respond and adapt to gradual temperature increases associated with climate change. The observed decline in meristem activity and overall root system architecture at high, but not lethal, temperatures highlights the need to define precise thresholds for non-lethal stress in plant development. From an agricultural viewpoint, such thresholds may represent hidden constraints that limit crop performance as temperatures rise.

In this study, we identified 31°C as a non-lethal thermal stress condition for *Arabidopsis* root development. Unlike lethal heat shock (above 37°C), which rapidly causes seedling death, exposure to 31°C allowed plants to survive but resulted in distinct developmental impairments. Non-lethal thermal stress reduced primary root growth, suppressed O₂^−^ accumulation, decreased root meristem size, and restricted PLT2 protein distribution at the seedling stage (Figs 1–3, 6). Although the overall structure of the apical meristem remained intact, cell division and progression within the meristematic zone markedly slowed down, leading to premature cell elongation and differentiation (Fig. 3). A transcriptome analysis of root developmental zones further showed the down-regulation of the genes related to cell proliferation and meristem maintenance instead of inducing canonical heat shock response genes (Figs 4b, c, d, S1). Seedlings could adapt and survive under non-lethal thermal stress but eventually developed shorter primary roots with shorter lateral roots (Figs 1, 7). Thus, while non-lethal thermal stress does not cause immediate death, its adaptive cost becomes evident as diminished growth and productivity (Figs 1, 7).

### Root developmental zone-specific transcriptomics as a breakthrough

Most previous transcriptomic studies on heat stress in *Arabidopsis* have primarily focused on the induction of classical stress-responsive genes under acute or moderately high temperature conditions (Larkindale & Vierling, 2008; Quint *et al*., 2016; Vu *et al*., 2019; Guo *et al*., 2022). Consequently, the majority of RNA-seq analyses have been performed using whole seedlings or entire roots, providing valuable global insights but lacking spatial resolution across distinct developmental zones. However, our approach targeted specific root zones at defined time points before and after the onset of root phenotype, allowing us to capture the earliest stages of adaptation to non-lethal heat stress.

This developmental zone- and time-resolved perspective reveals that roots respond to 31°C primarily through developmental signaling pathways (Fig. 4c, d), rather than by strongly activating canonical heat stress response genes (Fig. S1). Such findings highlight the importance of spatiotemporally resolved transcriptomics for uncovering novel mechanisms of plant adaptation to non-lethal environmental stress.

### Sensitivity of lateral roots to low concentrations of RGF2 and RGF8 under non-lethal thermal stress

A previous study showed that overexpression of RGF8 or treatment with high concentrations (2-4 µM) inhibits lateral root development under normal growth conditions (Fernandez *et al*., 2015). In contrast, our treatments used much lower concentrations (10-100 pM). These low concentrations had no effect under the control condition, but 50 pM RGF2 and RGF8 markedly restored lateral root elongation under non-lethal thermal stress (Fig. 7). However, 10 pM of these peptides had no effect, whereas 100 pM had negative effects. These observations suggest that RGF peptides possess an intrinsic activity threshold, and this threshold is lowered under non-lethal thermal stress, with the effective signaling range being set at extremely low concentrations around 50 pM. The molecular mechanism by which temperature shifts this signaling threshold remains unknown and will need to be elucidated in future studies.

### Topical application of RGF peptide stimulated a more complex root architecture under non-lethal thermal stress

Under non-lethal thermal stress, topical application of RGF peptides promoted not only primary root meristem maintenance but also lateral root growth, resulting in a more elaborate and integrated root system. Consistent with their distinct physiological roles, RGF1 primarily acted on the primary root meristem, whereas RGF8 mainly regulated lateral root development under non-lethal thermal stress (Figs 6, 7). Interestingly, RGF8 differs from RGF1 and RGF2 in that it cannot rescue the *tpst* mutant primary meristem defect (Matsuzaki *et al*., 2010), consistent with its specialized role in lateral root development. Together, these findings suggest that roots adapt to non-lethal thermal stress by differentially modulating primary- and lateral-root-associated RGF peptides.

The enhanced lateral root growth suggests that RGF signaling acts beyond the root tip, influencing post-meristematic developmental programs that shape overall root system architecture. These results suggest that a potential combined treatment of RGF1 and 8, by expanding both meristem activity and lateral growth, may more effectively counteract the inhibitory effects of elevated temperature and generate a more complex root structure capable of sustaining growth under stress.

### Potential of RGF peptide application to mitigate heat stress in crops

RGF1 and RGF8 peptides could serve as a potential tool to alleviate thermal stress without the need for genetic modification. Because these small peptides are stable and biologically active at even low concentrations (50 pM) (Fig. 7), topical application may provide a flexible and rapid means to enhance stress resilience in plants. RGF peptide treatment lowered H₂O₂ levels, which are typically elevated under various stress conditions, suggesting that this pathway could alleviate not only heat but also other abiotic stresses involving oxidative imbalance. Although peptide delivery through soil remains technically challenging, our successful application on agar medium indicates that this approach could be adapted to hydroponically cultivated crops such as rice and other water-grown species.

In our study, exogenous RGF1 treatment expanded the root meristem and conferred resistance to non-lethal thermal stress (31°C) (Fig. 6). In contrast, the roots of the *rgf1/2/3* and *rgfr1/2/3* mutants, which possess smaller meristems, were more susceptible to the thermal stress (Fig. 5). These results demonstrate that root meristem size is a critical determinant of the ability to withstand non-lethal thermal stress.

It is plausible that similar regulatory mechanisms function across diverse crops. *RGF* and *RGFR* genes are conserved in at least 16 species of angiosperms, including *Oryza sativa*, *Zea mays*, *Solanum lycopersicum*, and *Glycine max* (Fang *et al*., 2021). The mature peptide sequences are also highly conserved across species, and synthetic *Zea mays* RGF1 (ZmRGF1) peptide or overexpression of the *ZmRGF1* gene enlarges the meristematic zone in *Arabidopsis* roots (Fang *et al*., 2021). If the RGF-RGFR signaling pathway is both evolutionarily and functionally conserved, it may be possible to use *Arabidopsis* RGF peptide to alleviate abiotic stress in crops.

In this research, we demonstrated that RGF1, RGF2, and RGF8 peptides mitigated defects of primary root meristem and lateral root elongation under non-lethal thermal stress. In contrast, the *rgf1/2/3*, *rgfr1/2/3*, and *plt2* mutants showed hypersensitivity to the stress. Together, our findings highlight the potential of RGF peptide signaling as a natural, evolutionarily conserved mechanism that links developmental plasticity with environmental adaptation. Harnessing this pathway may provide a new foundation for enhancing plant resilience in the era of global climate change.

## Supporting information

Fig. 4_RNAseq_source_data

---

**Fig. S1.**
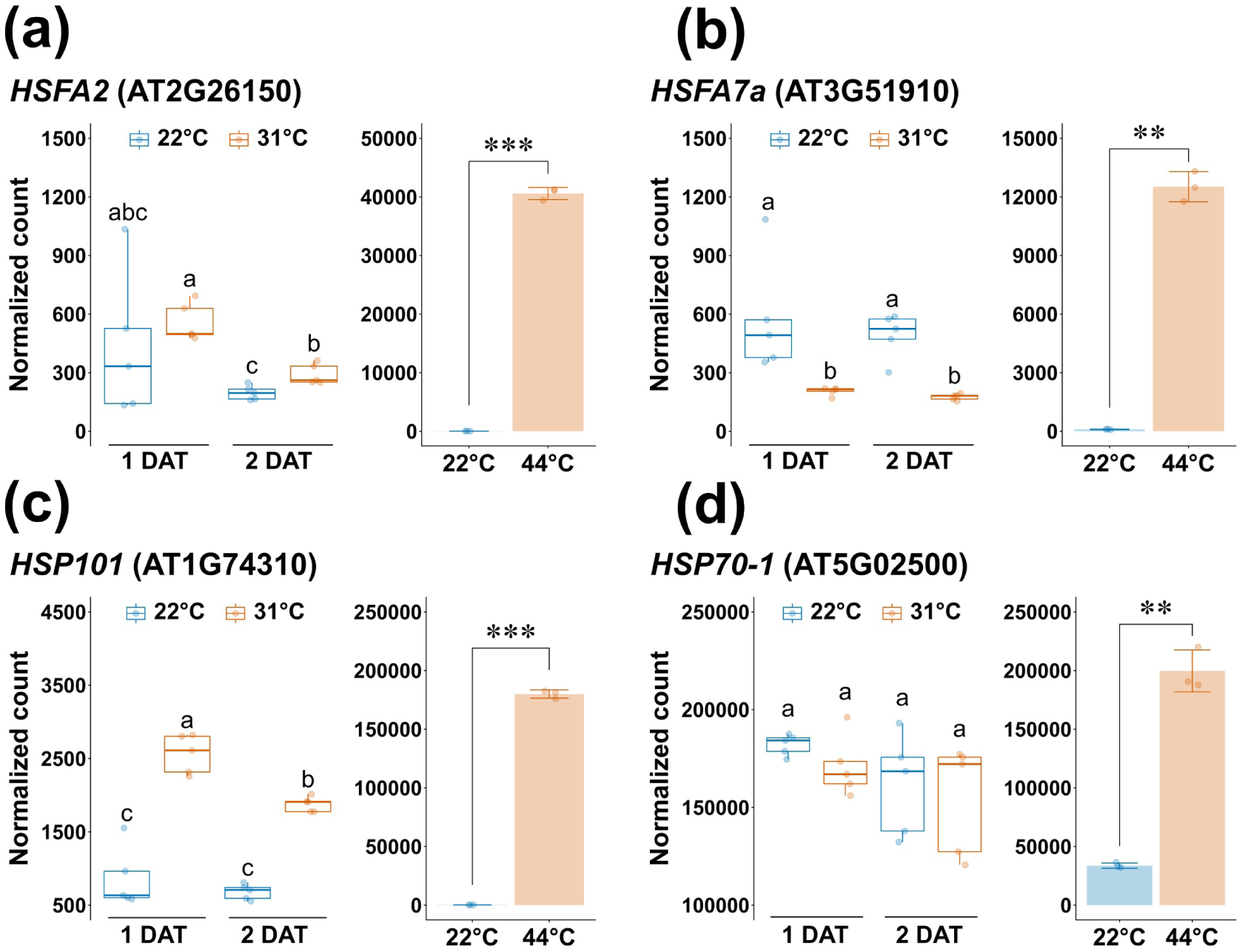
Non-lethal heat stress did not stimulate the expression of heat-shock responsive genes. Fig. S1 Non-lethal heat stress did not stimulate the expression of heat-shock responsive genes. (a‒d) Expression of heat stress marker genes. For the pairwise comparison of the data in this study (boxplots), different letters represent significant differences using ART-ANOVA + Games-Howell test (a), ART-ANOVA + Tukey-HSD test (b), and ANOVA + REGWQ test (c‒d). For the two-sample comparison of the data taken from Yang et al., 2023 (bar plots), independent t-test was conducted (a‒d). Asterisks indicate the gene expression of the heat treatment group is significantly different from the control (t-test; ***: p < 0.001, **: p < 0.01, *: p < 0.05), while ns indicates non-significant differences (p > 0.05). In the boxplots, the circles represent each individual data point, and sample sizes are shown below the corresponding boxes. The horizontal line within boxes represents the median, and the upper and lower hinges, respectively, indicate the 75th and 25th percentiles (interquartile range, IQR). The whiskers show the 1.5 extension of the IQR. In the bagplots, the circles represent each individual data point, and the error bar represent the standard deviation of the samples (n = 3).

## Acknowledgements

We thank Elizabeth Haswell, Kuo-Chen Yeh, Yee-Yung Charng, Su-Chiung Fang and Jian-You Wang for valuable comments on the manuscript; Elizabeth Haswell and Miranda Loney for English editing; Yuan-Fong Chin for assistance with root growth assays under non-lethal heat stress; the High Throughput Sequencing Core at the Biodiversity Research Center, Academia Sinica, for Illumina sequencing; the Confocal Microscope Core Facility at the Biotechnology Center in Southern Taiwan, Academia Sinica, for maintaining the confocal laser scanning microscope; and the NGS High Throughput Sequencing Core Facility, Academia Sinica, for maintaining the computing server.

This work was supported by a Career Development Award, Academia Sinica, Taiwan (AS-CDA-111-L05); the National Science and Technology Council, Taiwan (112-2311-B-001-021-MY3, 111-2311-B-001-028, 109-2313-B-001-006-MY3); and the Agricultural Biotechnology Research Center, Academia Sinica, Taiwan, to M.Y.

## Author contributions

Y-C.H. and M.Y. conceptualized the study; Y-C.H., S-Y.S., and M.Y. performed all experiments; J-K. L. performed the computational and statistical analyses; all the authors wrote the paper.

